# *Chlamydia trachomatis* TmeA promotes pedestal formation through N-WASP and TOCA-1 interactions

**DOI:** 10.1101/2024.10.31.621191

**Authors:** Alix McCullough, C.A. Jabeena, Brianna Steiert, Robert Faris, Mary M. Weber

## Abstract

*Chlamydia trachomatis* (*C.t.*) is the causative agent of several human diseases, including the sexually transmitted infection chlamydia and eye infection trachoma. As an obligate intracellular bacterial pathogen, invasion is essential for establishing infection and subsequent pathogenesis. To facilitate invasion, *C.t.* secretes effector proteins through its type III secretion system (T3SS). These effectors facilitate bacterial entry by manipulating multiple pathways involved in host actin cytoskeletal regulation. Previous studies have demonstrated that the T3SS effector protein TmeA is crucial for *C.t.* invasion, as it recruits and activates N-WASP. This interaction leads to recruitment and activation of the Arp2/3 complex, promoting cytoskeletal rearrangements at the invasion site to facilitate *C.t.* uptake. In this study, we define the role of the N-WASP CRIB domain in mediating this interaction, showing that TmeA acts as a functional mimic of Cdc42 in activating N-WASP. Additionally, we identified TOCA-1 as another host protein that directly interacts with TmeA. In other bacterial pathogens, notably Enterohemorrhagic *E. coli*, N-WASP and TOCA-1 are hijacked to mediate pedestal formation. Using siRNA to knockdown N-WASP and TOCA-1, followed by transmission electron microscopic, we observed that both N-WASP and TOCA-1 are important for in *C.t.*-mediated pedestal formation. Collectively, these findings reveal a unique mechanism of TmeA-mediated invasion, where direct interactions with N-WASP and TOCA-1 facilitate pedestal formation.

**Importance:** *Chlamydia trachomatis* (*C.t.*) is an obligate intracellular bacterial pathogen that poses a significant threat to human health, being associated with various diseases, including chlamydia— the most prevalent bacterial sexually transmitted infection—and trachoma. While Chlamydia infections are often asymptomatic, they can lead to serious complications such as sterility, ectopic pregnancy, and increased risk of cervical and ovarian cancers. Due to its intracellular nature, host cell invasion is essential for *C.t.* survival. Here, we present new data detailing the binding interactions between the *C.t.* invasion effector protein TmeA and host cell proteins N-WASP and TOCA-1, demonstrating that both N-WASP and TOCA-1 are involved in pedestal formation during *C.t.* invasion. This research advances our understanding of TmeA-mediated host cell invasion, illuminating a key pathway required for *C.t.*-mediated pathogenesis.

## Introduction

*Chlamydia trachomatis* (*C.t.*) is an obligate intracellular bacterial pathogen that causes a wide variety of diseases in humans, including trachoma and a sexually transmitted infection, chlamydia (1, 2). While *C.t.* infections are often asymptomatic in early stages, complications from infection can lead to scarring of the female genital tract, resulting in pelvic inflammatory disease, sterility, ectopic pregnancy, and an increased risk of cervical and ovarian cancers (3, 4). Reinfections are common due to lack of long-term immunity and the absence of a vaccine (5). Elucidating the molecular mechanisms underlying how *C.t.* invades epithelial cells and causes disease is vital for the development of improved therapeutics and identification of vaccine candidates.

Because *C.t.* is an obligate intracellular pathogen, invasion is a critical step in its lifecycle. As such, it has developed multiple mechanisms to invade host cells. At the end of its developmental cycle, *C.t.* prepackages a subset of type III secreted (T3S) effector proteins, including invasion-specific effector proteins TarP (<u>T</u>ranslocated <u>a</u>ctin recruiting <u>P</u>hosphoprotein), TepP (<u>T</u>ranslocated <u>e</u>arly <u>p</u>hospho-<u>P</u>rotein), TmeA (<u>T</u>ranslocated <u>m</u>embrane <u>e</u>ffector <u>A</u>), and TmeB (<u>T</u>ranslocated <u>m</u>embrane <u>e</u>ffector <u>B</u>) (6–10). Tarp binds F- and G-actin directly to facilitate actin filament bundling and elongation, as well as interacting with host effectors to stimulate Rac1 signaling pathways and actin branching through the Arp2/3 complex (11–15). Notably, TarP has been shown to localize to *C.t.*-associated actin pedestals, though its role in pedestal generation has not been explored (6). TmeB has been shown to inhibit the Arp2/3 complex, suggesting it may have a part in disassembly of TmeA- and/or TarP-generated actin structures during invasion (16). TepP plays a role in modulating the innate immune response during early infection, including dampening of type I IFN responses via recruitment and activation of signaling adaptor CrkL and class I phosphoinositide 3-kinases (PI3K) to nascent inclusion membranes, leading to generation of phosphoinositide-(3,4,5)-triphosphate (PIP3) (8, 17). TepP expression additionally reduces neutrophil recruitment in organoid models and disrupts epithelial cell-cell junctions through interactions with EPS8 (18, 19).

TmeA has been shown to interact with the host protein AHNAK; however, the function of this interaction during infection remains to be fully elucidated and is distinct from its role in invasion (20). TmeA additionally recruits and activates host N-WASP (<u>N</u>eural <u>W</u>iskott-<u>A</u>ldrich <u>s</u>yndrome <u>p</u>rotein) and thus the Arp2/3 complex, resulting in actin remodeling (21, 22). TmeA-N-WASP interactions stimulate uptake of *C.t.* independent of TarP, but it also contributes to a TarP-mediated invasion pathway through oligomerization of host dynamin-2 (Dyn2) (23). N-WASP is modulated by multiple bacterial pathogens, including *Chlamydia pneumoniae*, *Salmonella enterica* Typhimurium, *Brucella abortus*, *Shigella flexneri*, and Enterohemorrhagic *Escherichia coli* (EHEC) (24–29). N-WASP is an autoinhibited protein, meaning its <u>G</u>TPase <u>b</u>inding <u>d</u>omain (GBD) binds to and inhibits the activity of the C-terminal VCA domain (30). Upon binding of small GTPases, primarily Cdc42, to the CRIB motif of the GBD, N-WASP is relieved of autoinhibition, exposing the VCA domain. This exposure enables N-WASP to bind to the Arp2/3 complex, facilitating Arp2/3-mediated actin branching and ultimately leading to the formation of host cell structures including filopodia and *E. coli* pedestals (28–33). N-WASP forms complexes with additional host proteins including <u>W</u>ASP-Interacting <u>P</u>rotein (WIP) and <u>T</u>ransducer <u>O</u>f <u>C</u>dc42-dependent <u>A</u>ctin Assembly (TOCA-1), which modulate its activity (34, 35). WIP regulates N-WASP activity and helps to stabilize actin filaments, while TOCA-1 works in conjunction with Cdc42 to help activate the N-WASP/WIP complex (34, 35).

EHEC uses a T3S effector, EspF_u,_ to recruit and activate N-WASP to facilitate pedestal formation (28, 29). Notably, it additionally recruits and activates TOCA-1, which is important for efficient EHEC pedestal formation (31). *S. flexneri* also recruits TOCA-1 to its actin cocoons and actin (36, 37). Based on TmeA’s role in N-WASP recruitment and modulation, as well as the prior identification of *C.t.*-associated pedestals, we hypothesized TOCA-1 may also play a role in *C.t.* (6, 21, 22, 38). Herein, we further characterize the TmeA-N-WASP interaction, showing that TmeA binds N-WASP via the N-WASP CRIB motif. We additionally identify TOCA-1 as a host protein that is bound directly by TmeA, independent of TmeA-N-WASP interactions. Lastly, we show a role for both proteins in *C.t.*-induced pedestal formation.

## Results

### TmeA binds to the Cdc42 binding site of N-WASP

Our previous study found that the GTPase Binding Domain (GBD) ligand motif of TmeA was important for TmeA-N-WASP binding, and Keb *et al*. has shown that TmeA is able to directly activate N-WASP (21, 22). However, TmeA’s specific binding site on N-WASP has not been determined. Because the GBD ligand motif of TmeA was important for the interaction, and this ligand motif is utilized by EspF_u_ to bind the autoinhibitory VCA binding site of the N-WASP GBD, we hypothesized that TmeA would similarly bind the VCA binding site of N-WASP. To test this hypothesis, we first designed FLAG-tagged N-WASP deletion constructs (Fig 1A) of the entire GBD (ΔGBD, aa191-275), the region of the GBD containing the Cdc42 binding site (ΔCdc42, aa191-237), or the region of the GBD containing the autoinhibitory VCA domain binding site (ΔVCA, aa215-275) (Fig 1A). We co-transfected cells with these plasmids in conjunction with GFP-tagged TmeA, or TmeB as a negative control, and immunoprecipitated the FLAG-tagged constructs. Using this approach, we surprisingly found that the Cdc42 binding region, but not the VCA binding region, was necessary for TmeA-N-WASP interaction (Fig 1B).

**Figure 1:**
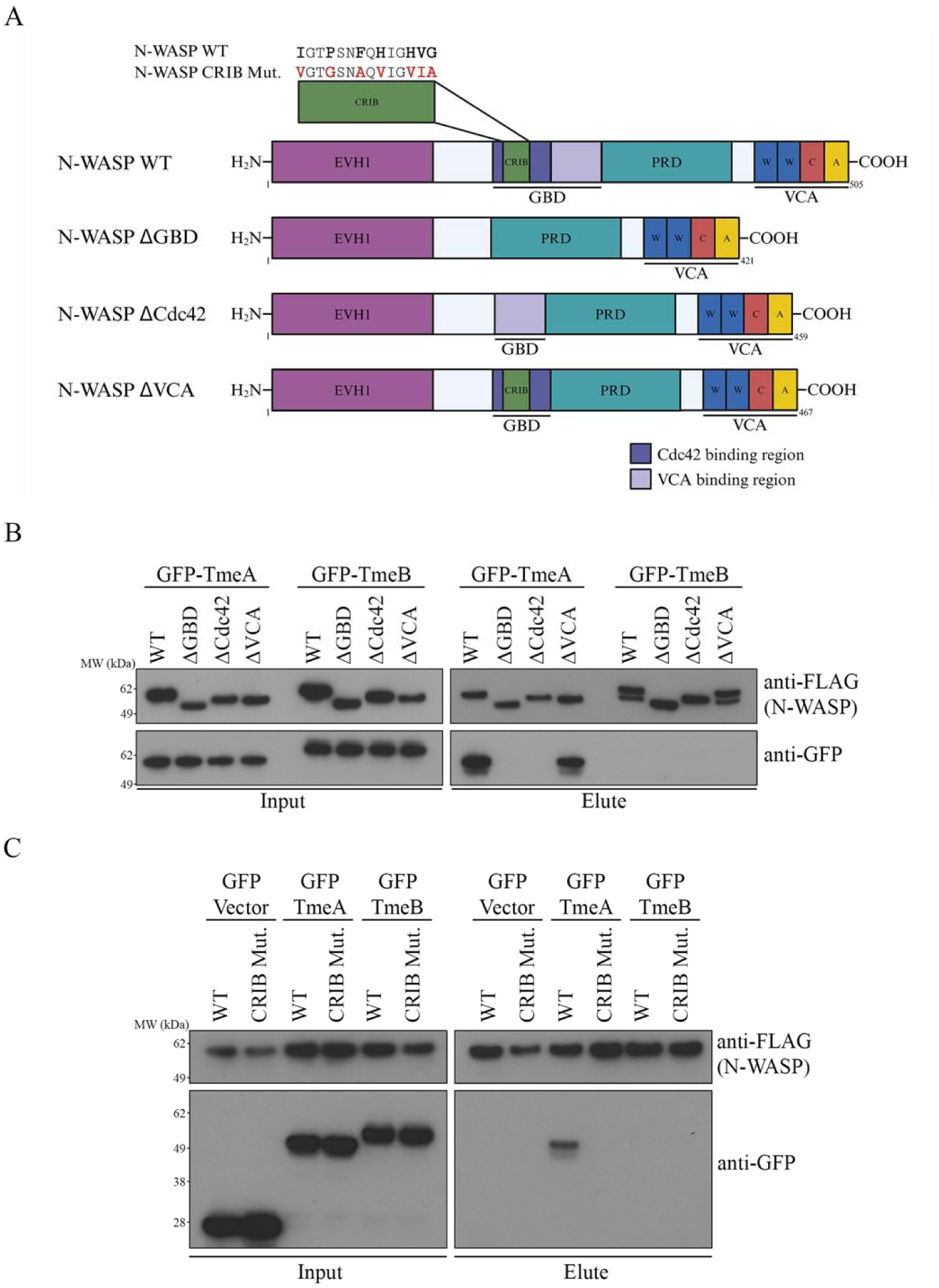
TmeA binds to the Cdc42 binding site of N-WASP. **A.** Schematic depicting N-WASP deletion constructs. **B,C.** FLAG-tagged N-WASP constructs were co-transfected with GFP-tagged C.t. effectors in HeLa cells. The FLAG tagged proteins were immunoprecipitated and samples were probed with anti-GFP or anti-FLAG antibodies.

To investigate this interaction further and determine whether TmeA binds at the same site as Cdc42, we designed conservative amino acid replacements critical for the N-WASP CRIB motif (**I**GT**P**SN**F**Q**H**IG**HVG** → **V**G**TG**SN**A**Q**V**IG**VIA**) We then performed a co-immunoprecipitation as described above. This motif is found in various Cdc42/Rac effectors including N-WASP and is necessary, though not sufficient, for Cdc42 binding (39). Here, we demonstrate that the CRIB motif is necessary for TmeA-N-WASP interactions, as shown by the loss of TmeA co-IP with the N-WASP CRIB mutant construct (Fig 1C). These results confirm that TmeA binds to N-WASP as a functional mimic at the same site as Cdc42 during infection.

### TmeA directly binds TOCA-1, independent of N-WASP and Cdc42 binding sites

Given TOCA-1’s role in N-WASP activation and its recruitment to EHEC pedestals as well as to *Shigella flexneri* actin tails and cocoons alongside N-WASP (28, 29, 31, 36, 37), we hypothesized that TmeA might also interact with TOCA-1. To test this, we first transfected GFP-TmeA, a TmeA GBD ligand motif mutant deficient in N-WASP binding (TmeA GBD) (21), TmeB, or GFP vector into HeLa cells and immunoprecipitated them before probing for endogenous TOCA-1 (Fig 2A). TmeA and the TmeA GBD construct were both able to co-IP with TOCA-1, indicating that TmeA interacts with TOCA-1 independent of N-WASP interactions.

**Figure 2:**
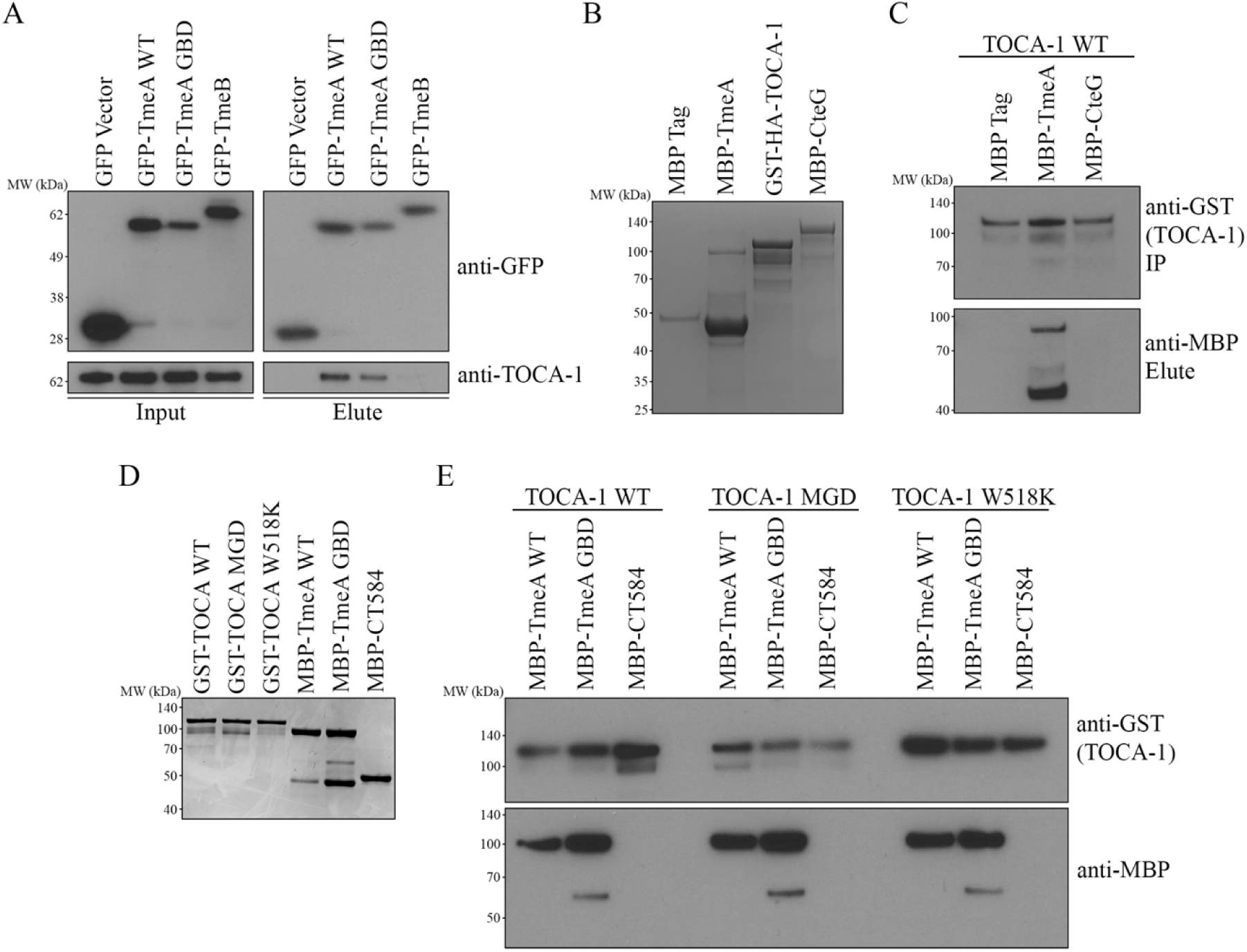
TmeA directly binds TOCA-1 independent of Cdc42 and N-WASP binding sites. **A.** GFP-tagged C.t. effectors were transfected in HeLa cells. The GFP tagged proteins were immunoprecipitated and samples were probed with anti-GFP and anti-TOCA-1 antibodies. **B,D.** GST-tagged TOCA-1 constructs and MBP-tagged C.t. effectors were expressed in E. coli and purified using GST or MBP resin on gravity columns. Recombinant protein expression was confirmed with Coomassie staining. **C, E.** Recombinant GST-tagged TOCA-1 constructs were bound on GST resin. Bound proteins were incubated with MBP-tagged C.t. effectors and eluted, and samples were probed with anti-GST and anti-MBP antibodies.

We thus hypothesized that similar to EspF_u_, TmeA may directly bind to TOCA-1. To test this, we expressed and purified MBP-TmeA, MBP-CteG, MBP tag, and GST-TOCA-1 (Fig 2B). We performed a recombinant GST pulldown of the MBP tagged proteins. We utilized CteG, an effector previously shown to bind to CETN2 (40), as a negative control for non-specific binding interactions. We found that TOCA-1 specifically and directly bound to TmeA (Fig 2C), emphasizing this is a direct interaction between TmeA and TOCA-1 that is independent of other host protein complexes. Because TmeA bound to TOCA-1 independent of both its GBD ligand motif and N-WASP, we wanted to further test whether known Cdc42 or SH3 binding sites on TOCA-1 may play a role in direct TmeA-TOCA-1 interactions. TOCA-1 binds Cdc42 through its HR1 domain and interacts with N-WASP via its SH3 domain (35). We thus performed a co-IP with TOCA-1 constructs, TmeA, TmeA GBD, or CT584 as a negative *C.t.* effector control (41). The TOCA-1 constructs were TOCA-1 WT, a TOCA-1 MGD mutant, which lacks GTPase binding activity, and a W518K mutant, which lacks SH3 binding activity (35) (Fig 2D). Both mutants were able to co-IP with TmeA WT and TmeA GBD, showing that neither the Cdc42 nor N-WASP binding sites are targeted as essential residues by TmeA (Fig 2E). Taken together, we have shown that TOCA-1 is a direct interactor of TmeA independent of N-WASP or other host proteins, and this interaction does not use TOCA-1 canonical binding motifs.

### N-WASP and TOCA-1 play a role in *C.t.* pedestal formation

*C.t.* has previously been shown to form pedestals during invasion of host cells (6, 38). Because of TOCA-1’s role in EspF_u_-mediated pedestal formation, we hypothesized that TOCA-1 may play a similar role in *C.t.* pedestal formation during invasion. To test this, we knocked down expression of TOCA-1 and N-WASP using siRNA, and then challenged KD or mock KD control HeLa cells with WT L2 *C.t.* for 15 minutes (Fig. 3A). We used transmission electron microscopy (TEM) to evaluate structures associated with EBs at the cell surface (Fig. 3B, Sup. Fig. 1). Compared to the mock control, EBs on both N-WASP and TOCA-1 KD cells displayed a significant reduction in association with pedestals (Fig 3C), implying that both N-WASP and TOCA-1 play a role in *C.t.-*mediated pedestal formation.

**Figure 3:**
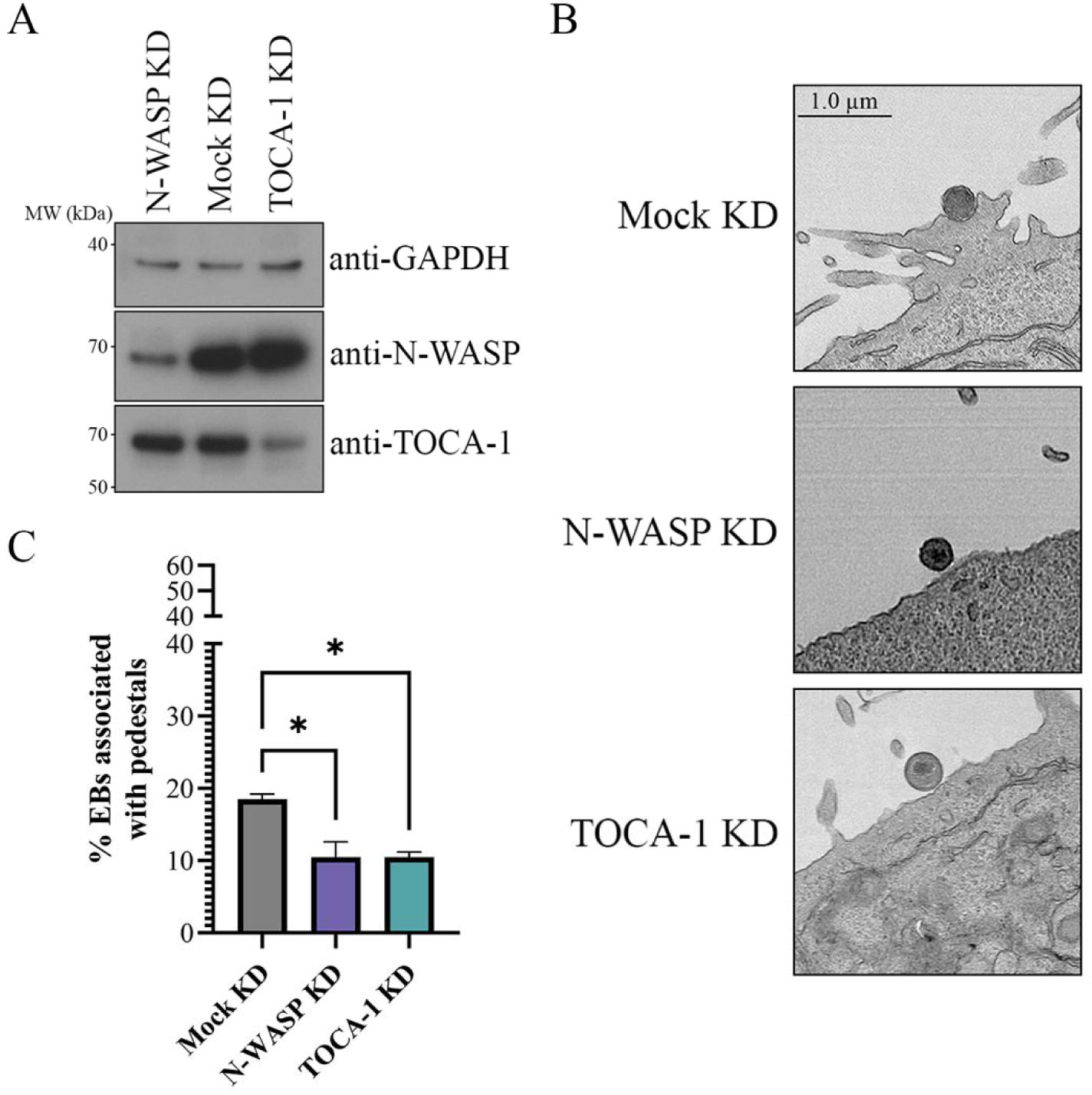
N-WASP and TOCA-1 play a role in C.t. pedestal formation. **A.** N-WASP and TOCA-1 were knockdowned in HeLa cells. Knockdown was verified using western blotting, probing with anti-GAPDH, anti-N-WASP, and anti-TOCA-1 antibodies. **B.** HeLa cells were asynchronously infected with WTL2 at an MOI of 50 for 15 minutes and imaged with transmission electron microscopy, three representative images are shown. **C.** Quantification of EBs associated with pedestals. 100 EBs per experiment were assessed from two separate experiments, in which images were blinded and categorized as associated or not associated with pedestals. EBs associated with pedestals were compared to total EBs to determine the percentage associated with pedestals.

## Discussion

TmeA is an important T3S effector protein that appears to interact with several cytoskeletal proteins to promote *C.t.* infection. Interactions with the host protein AHNAK function to inhibit actin bundling, although this interaction is not important for *C.t.* mediated invasion and its role during *C.t.* infection remains unknown (20). During the invasion phase, TmeA interacts with N-WASP, recruiting it to the invasion site to activate the Arp2/3 complex to promote actin branching (21, 22). Given the compelling evidence for TmeA’s ability to directly activate N-WASP (22), we set out to identify the precise binding site on N-WASP to gain deeper mechanistic insight into this interaction. Other bacterial effectors, such as *Shigella flexneri* IcsA and Enterohemorrhagic *E. coli* EspF_u_, have been shown to bind to the VCA binding region of the N-WASP GBD, while *Chlamydia pneumoniae* SemD acts as a molecular mimic of Cdc42 and binds the N-WASP CRIB domain (24, 29, 42). Prior work identified a GBD ligand motif in TmeA that is essential for co-immunoprecipitation with N-WASP (21). Furthermore, TmeA was shown to be sufficient for N-WASP activation *in vitro* (22). These two key observations lead us to hypothesize that TmeA likely binds the VCA binding region of the N-WASP GBD during *C.t.* invasion. Our new data oppose this hypothesis and rather indicate that, like SemD, the N-WASP CRIB motif is necessary for TmeA co-IP and that the VCA binding region of the GBD is dispensable for these interactions. This indicates that TmeA is activating N-WASP by binding at the Cdc42 binding site, likely acting as a molecular mimic to directly activate N-WASP (Fig. 4). These data are supported by prior evidence that Rac, but not Cdc42, was activated by and recruited to the site of *C.t.* invasion (43). However, other studies have shown that Cdc42 does appear to be recruited to the site of invasion and plays a role in TmeA-mediated invasion, as its knockdown leads to an invasion defect in *C.t.* that express TmeA (22, 44). Further study is required to determine what role Cdc42 may play in TmeA-mediated invasion.

**Figure 4:**
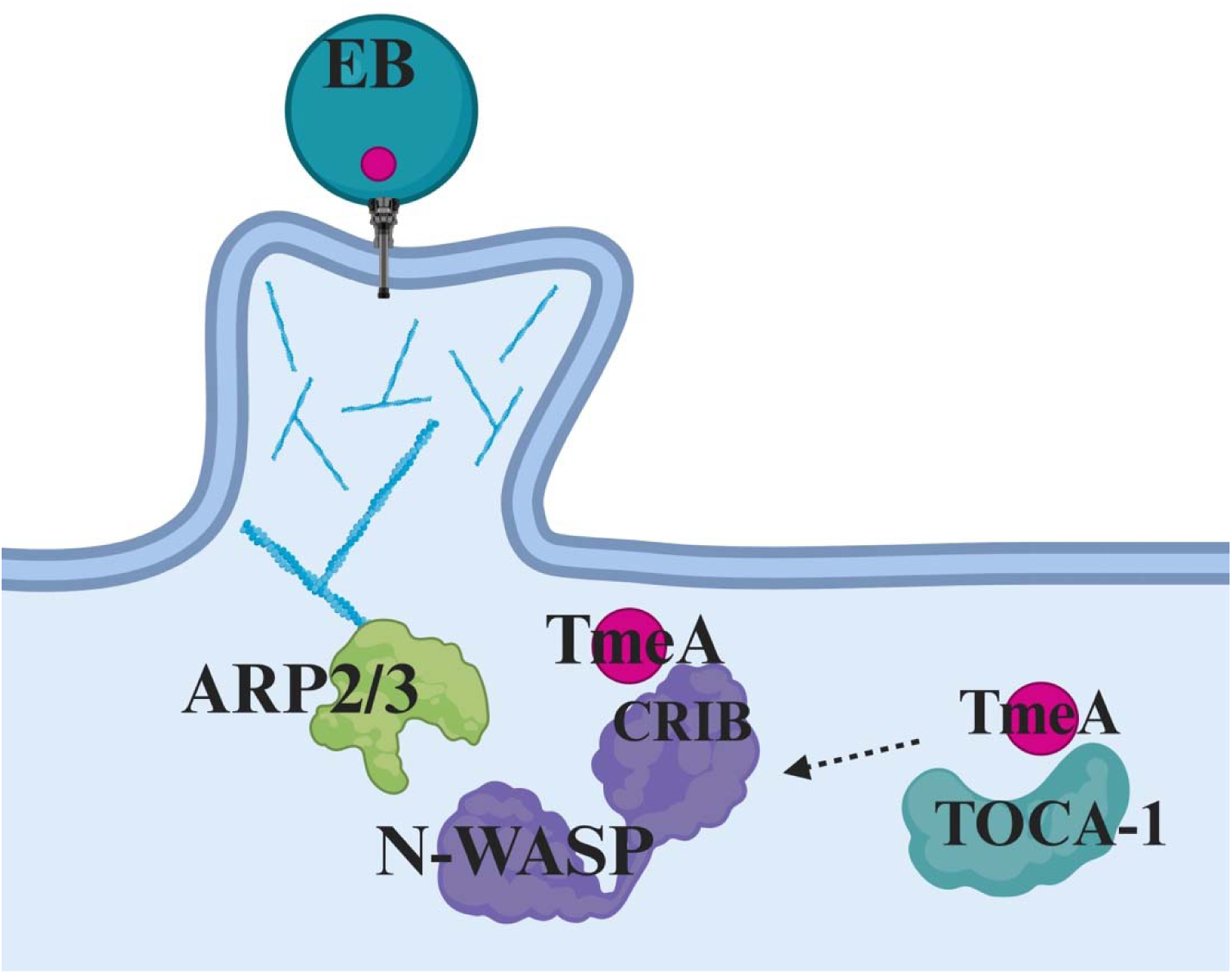
TmeA interacts with N-WASP and TOCA-1, which leads to EB-pedestal association. TmeA interacts with N-WASP via the N-WASP CRIB domain to activate it as a Cdc42 molecular mimic. It additionally interacts with TOCA-1 independently of N-WASP. The convergent roles of N-WASP and TOCA-1 in EB pedestal association indicate that these interactions likely contribute to a TmeA-mediated invasion pathway where EBs are taken up via pedestals.

TOCA-1 has also been implicated as an interacting partner of N-WASP. Notably, it is targeted by several bacterial effector proteins (31, 36, 37). TOCA-1-N-WASP interactions play a role in endocytic membrane trafficking (45, 46). The F-BAR domain of TOCA-1 binds to membranes and induces membrane curvature, as well as recruits N-WASP to the membrane (47). TOCA-1 binds Cdc42 through its HR1 domain and interacts with N-WASP via its SH3 domain (35). Analysis of binding dynamics reveals that Cdc42 exhibits a significantly higher affinity for N-WASP than for TOCA-1, suggesting that TOCA-1 may facilitate N-WASP activation by “handing off” Cdc42 to N-WASP (48). Notably, these studies were conducted using the purified TOCA-1 HR1 domain alone and a tertiary complex between TOCA-1 HR1, N-WASP GBD, and Cdc42 was not observed. However, evidence from Förster resonance energy transfer (FRET) studies suggest potential tertiary complex formation, which may lead to stabilization of the flexible proline rich domain (PRD) of N-WASP by the TOCA-1 SH3 domain (35, 46).

Both TOCA-1 and N-WASP have previously been linked to critical cellular processes including endocytosis (45, 46), EHEC pedestal formation (31), and *Shigella flexneri* actin cocoons (36, 37). Given these roles, we explored whether TOCA-1 similarly contributes to *C.t.* infection. Our findings indicate that TmeA can bind TOCA-1, and that these interactions are not abrogated by TOCA-1 mutants deficient in Cdc42 or N-WASP interactions with either ectopically expressed or recombinant proteins. We were unable to generate a TmeA truncation mutant deficient in TOCA-1 binding (Sup. Fig. 2), suggesting that TmeA might bind to multiple binding sites on TOCA-1. This hypothesis is supported by Alphafold 2.0 and HDOCK modeling, which predicts interactions at multiple sites on both proteins (Sup. Fig. 2). Future studies will determine whether TmeA can bind to TOCA-1 and N-WASP simultaneously and whether it exhibits differing affinities for either protein, similar to Cdc42.

Our data suggest a role for N-WASP and TOCA-1 in pedestal formation as knockdown of either protein significantly reduced EB-associated pedestals. Interestingly, TarP has previously been implicated in *C.t.* associated pedestals (6), thus, investigating whether TmeA and TarP co-contribute to the formation of these structures represents an intriguing area for future research. New work proposes a model of Dyn2 dependent *C.t.* invasion, where TarP recruits and TmeA activates Dyn2 (23). Dyn2 has notably been implicated in enteropathogenic *E. coli* pedestal formation alongside N-WASP (49). Thus, it is interesting to speculate that *C.t.* pedestals form via a collaborative pathway between TmeA and TarP.

In *E. coli*, actin pedestals are not utilized for invasion of host cells but rather to facilitate colonization via attachment to the intestinal mucosa (50). There is evidence that this attachment is important not only for anchoring the pathogen to the epithelium to prevent flushing from the colon, but also to allow for actin-based motility and cell-to-cell spread of EPEC and EHEC (50–52). It may additionally allow for more efficient translocation of secreted effectors into the host cell, although this could be due to attachment rather than specifically pedestal formation (53, 54). Notably, *C.t.* pedestal formation appears to lead to host cell internalization, whereas *E. coli* pedestal formation does not, highlighting an interesting area of future research to elucidate the differences between the two pathogens.

Taken together, we have shown that TmeA relies on the CRIB domain of N-WASP for co-IP, potentially functioning as a molecular mimic of Cdc42. Additionally, we have shown that TmeA directly interacts with TOCA-1 independent of its interactions with N-WASP. These interactions are crucial for *C.t.-*mediated pedestal formation, as knockdown of either host protein leads to a significant reduction in EB-pedestal association (Fig. 4). Overall, this work adds to the growing body of knowledge surrounding *C.t.* invasion mechanisms.

## Materials and Methods

### Bacterial and Cell Culture

Wild type and *tmeA-lx* (55) *Chlamydia trachomatis* serovar L2 (LGV 434/Bu) was propagated in HeLa 229 cells (American Type Culture Collection). Purification of EBs was performed using a Gastrografin gradient as previously described (56). HeLa cells were cultured at 37°C under 5% CO_2_ in RPMI 1640 supplemented with L-Glutamine, 10% Fetal Bovine Serum (FBS) (26140079, ThermoFisher Scientific), sodium bicarbonate (25080094, ThermoFisher Scientific), sodium pyruvate (11360070, ThermoFisher Scientific), and 50 µg/ml gentamicin (15750078, ThermoFisher Scientific).

### Cloning

For ectopic expression, TmeA, TmeA GBD (21), and TmeB were cloned into pcDNA3.1+N-eGFP (Genscript) using KpnI or NotI/XbaI sites. TmeA GBD truncations were generated from the TmeA GBD construct and cloned into pcDNA3.1+N-eGFP (Genscript) using Kpn/Not sites. pcDNA 3.1 FLAG N-WASPΔGBD (21), N-WASPΔCdc42, N-WASPΔVCA, and N-WASP CRIB mutant were generated by GenScript. pCS2+MT-hTOCA-1 WT, MGD, and W518K (Addgene 33030, 33033, and 33031) were cloned into pcDNA 3.1 HA (GenScript) using KpnI/XhoI sites. For recombinant protein expression, CteG, TmeA, TmeA GBD, and CT584 were cloned into pMAL-c5VT (University of Iowa Protein and Crystallography Facility) using NotI/SalI sites. Codon optimized pGEX 6P1 TOCA-1 WT, MGD, and W528K were purchased from GenScript.

### Co-Immunoprecipitation

HeLa cells were seeded at 4×10^5^ in 6-well plates (10062-892, VWR) 24 hours prior to transfection. GFP-tagged TmeA, TmeA GBD, TmeB, or empty vector were transfected alone or co-transfected with FLAG-tagged N-WASP WT, N-WASPΔGBD (21), N-WASPΔCdc42, N-WASPΔVCA, or N-WASP CRIB mutant. GFP-tagged TmeA GBD truncations were co-transfected with HA-tagged TOCA-1 WT. HeLa cells were transfected using Lipofectamine LTX (15338100, ThermoFisher Scientific) per the manufacturer’s instructions. At 18 hours post-transfection, cells were washed with 1x PBS (10010023, Gibco) and lysed using Eukaryotic Lysis Solution (ELS) (50mM Tris HCl, pH7.5 (15567-027, Invitrogen), 150mM NaCl (S23020, RPI), 1mM EDTA (E57020, RPI), and 1% Triton-X 100 (BP151-500, ThermoFisher Scientific) with Halt protease inhibitor cocktail (78430, ThermoFisher Scientific). Lysates were incubated on ice for 20 minutes, followed by pelleting of cell debris via centrifugation at 12,000xg for 20 minutes at 4°C. Cell free supernatants were applied to anti-GFP mAb-Magnetic Beads (D153-11, MLB) for 2 hours at 4°C or anti-FLAG M2 Affinity Gel (A2220, Sigma) overnight at 4°C. Unbound proteins were removed by washing in ELS without Triton-X 100 and the GFP or FLAG tagged proteins were eluted using NuPAGE LDS Sample Buffer (NP0007, ThermoFisher Scientific) and heated at 100°C for 5min. Samples were analyzed by western blotting.

### Western Blotting

Supernatants and eluted samples were run on 4-12% SurePAGE™ Bis-Tris Gels (M00652, M00653, or M00654, GenScript), followed by wet transfer to Immobilon-P polyvinylidene difluoride (PVDF) membranes (IPVH0010, Sigma-Aldrich). Membranes were probed with primary antibodies against TOCA-1 (1:4000, PA5-85726, Invitrogen), GFP (1:10000, NB600-308, Novus Biologicals), FLAG (1:4000, 701629, Invitrogen), or HA (1:4000, H6908, Sigma-Aldrich) followed by HRP-conjugated secondary antibodies. (1:10000; Rabbit: 1706515, BioRad; Mouse: 31430, ThermoFisher Scientific), then detected using ECL Prime Western Blotting Detection Reagent (RPN2236, Sigma Aldrich) and X-ray film.

### Recombinant Protein Purification

GST (glutathione S-transferase) Vector (pGEX6p1), MBP (maltose-binding protein) Vector (pMALc5V2), MBP-TmeA, MBP-TmeA GBD, MBP-CteG, and MBP-CT584 were expressed in BL21 (DE3) *E. coli* and codon-optimized GST-TOCA-1 WT, GST-TOCA-1 MGD, and GST-TOCA-1 W518K were expressed in Rosetta (DE3) *E. coli* (70954, Novagen). Transformants were inoculated in 50 ml of Luria Broth (LB) (L24400, RPI), then incubated at 37°C overnight. For expression of MBP-tagged proteins, glucose (G32045, RPI) was added to LB broth at a concentration of 2g/L. Overnight cultures were added to 950 ml LB broth and grown to OD 0.6-0.8. Protein expression was induced with 1mM Isopropyl-β-D-thiogalactopyranoside (IPTG) (I56000-25.0, RPI), followed by overnight induction at 18°C. *E. coli* was lysed in MBP (20mM Tris HCl pH7.5, 200mM NaCl, 1mM EDTA, 1 mM sodium azide (RTC000068, Sigma Aldrich), 10% glycerol (BP229-1, ThermoFisher Scientific), 1mM dithiothreltol (DTT) (D11000, RPI)) or GST (50mM Tris HCl pH7.5, 500 mM NaCl, 10% glycerol, 1mM DTT) column/lysis buffer via sonication with 1 pulse ON for 1 second and 1 pulse OFF for 1 second for 3 minutes at 70% amplitude. Lysed cells were pelleted at 10,000xg for 30 minutes at 4°C and supernatants were collected. 1 ml Pierce™ Glutathione Agarose (16100, ThermoFisher Scientific) or amylose resin (E8021S, NEB) for GST or MBP tagged proteins, respectively, was added to 25 ml gravity columns and washed with the appropriate column/lysis buffer. Supernatants were incubated in columns at 4°C for 1 hour for protein conjugation. The columns were then washed with 75 ml lysis/column buffer before addition of 2 ml elution buffer for GST-tagged (50mM Tris HCl, pH7.5, 150mM NaCl, 10mM L-glutathione reduced (G22010, RPI)) or MBP-tagged (10mM D-(+)-maltose monohydrate (M22000, RPI)) proteins. Columns were incubated with rotation for 30 minutes at 4°C and elutions were collected in 1 ml fractions, with a total of 4 fractions after 2 elutions. Protein concentration was determined via A280 measurement on a Nanodrop 2000 (ThermoFisher Scientific) and protein purity was confirmed via Coomassie staining (1610436, Bio-Rad) of proteins on SDS-PAGE gels.

### Recombinant Protein Pull-down

Buffer exchange was performed on GST-tagged proteins (GST-TOCA-1 WT, GST-TOCA-1 MGD, GST-TOCA-1 W518K) and MBP-tagged proteins (MBP-TmeA, MBP-TmeA GBD, MBP-CT584) using Amicon™ Ultra Centrifugal Filter Units (UFC901024, Millipore) to remove reduced glutathione. Proteins were added to the filters with 10 ml GST column/lysis buffer or MBP column buffer, followed by 4°C centrifugation at 4,000xg for 15 minutes, for a total of five centrifugations. Protein concentration after buffer exchange was determined via A280 measurement on a Nanodrop 2000 (ThermoFisher Scientific). 40µg of the GST-or MBP-tagged bait proteins were conjugated to 0.5 ml Pierce™ Glutathione Agarose (ThermoFisher Scientific) or amylose resin (NEB), respectively, as described above (Recombinant Protein Purification), followed by three washes with appropriate column/lysis buffer. 40µg of the prey proteins were added to the columns and incubated overnight at 4°C with rotation. Columns were washed three times with GST column/lysis buffer and proteins were eluted as described above (Recombinant Protein Purification) (40). Eluted proteins were analyzed by western blotting. Membranes were probed with primary antibodies against MBP (1:4000, R29.6, Santa Cruz Biotech) or GST (1:4000, 8-326, Invitrogen).

### Transmission Electron Microscopy

HeLa cells were seeded at 2.5×10^5^ in 6-well plates 24 hours prior to siRNA knockdown. *FNBP1L* (TOCA-1) or *WASL* (N-WASP) expression was knocked down using DharmaFECT 1(T-2001-03, Horizon Discovery) with SmartPool siRNA (*FNBP1L* 020718-01-0020, *WASL* L-006444-00-0020, Horizon Discovery) according to the manufacturer’s protocol (Horizon Discovery). 48 hours post-transfection, knockdown was verified by western blotting using primary antibodies, all at a concentration of 1:4000, against TOCA-1 (PA5-85726, Invitrogen), N-WASP (NBP1-82512, Novus Biologicals), or a GAPDH loading control (SAB4300645, Sigma-Aldrich). Simultaneous to knockdown verification, remaining cells were asynchronously infected with WT L2 *C.t.* at an MOI of 50 for 15 minutes. Cells were fixed using 2.5% glutaraldehyde at 4°C overnight (21). Cells were post-fixed using 1% Osmium Tetroxide and 1.5% Potassium Ferrocyanide, followed by staining using 2.5% Uranyl Acetate. Samples were dehydrated using increasing concentrations of EtOH (50-100%) before embedding in Eponate12 resin (Ted Pella, Inc.). Using a Leica UC6 ultramicrotome (Leica Microsystems), 80 nm thin sections were made (21). Images were taken on a Hitachi HT7800 transmission electron microscope with an AMT NanoSprint15 high-resolution, high-sensitivity camera system. Two replicates of 100 EBs per sample were imaged, blinded, and scored for association with pedestals, filopodia-like structures, pits, or no cell structure utilizing example images of the structures from *Faris et al.* (2019), *Carabeo et al.* (2002), and *Clifton et al.* (2004).

### Molecular docking of TmeA and TOCA-1

The tertiary structure of TmeA and Toca-1 was predicted using AlphaFold 2.0 (57). The generated structures were used for protein docking using Hdock server (58). The resultant docking file gave possible interface residue pairs between TmeA and TOCA-1 and a confidence score. Binding is very likely when the confidence score is above 0.7.

### Statistical Analysis

One-way ANOVAs were used with Tukey’s multiple comparisons post-test, with *P* < 0.05 (*). Analysis was performed using GraphPad Prism 9.4.1 software

## Acknowledgments

We acknowledge grant support from the NIH (M.M.W., R01 AI150812, R01 AI155434, and R61 AI179999; A.M., T32 AI007511, B.S. T32 AI007511) and the University of Iowa Stead Family Scholars to M.M.W. We thank Weber lab members Steve Huang, Paige McCaslin, and Parker Smith for assistance with data analysis, as well as Tom Moninger from the University of Iowa Central Microscopy Research Facility for training in ultramicrotomy and TEM. We additionally thank the University of Iowa Central Microscopy Research Facility (CMRF) for use of the Leica EM UC6 Ultramicrotome and Hitachi HT7800 TEM. We kindly thank Ken Fields for sharing the *tmeA*::*lx* strain.

---

**Supplemental Figure 1.**
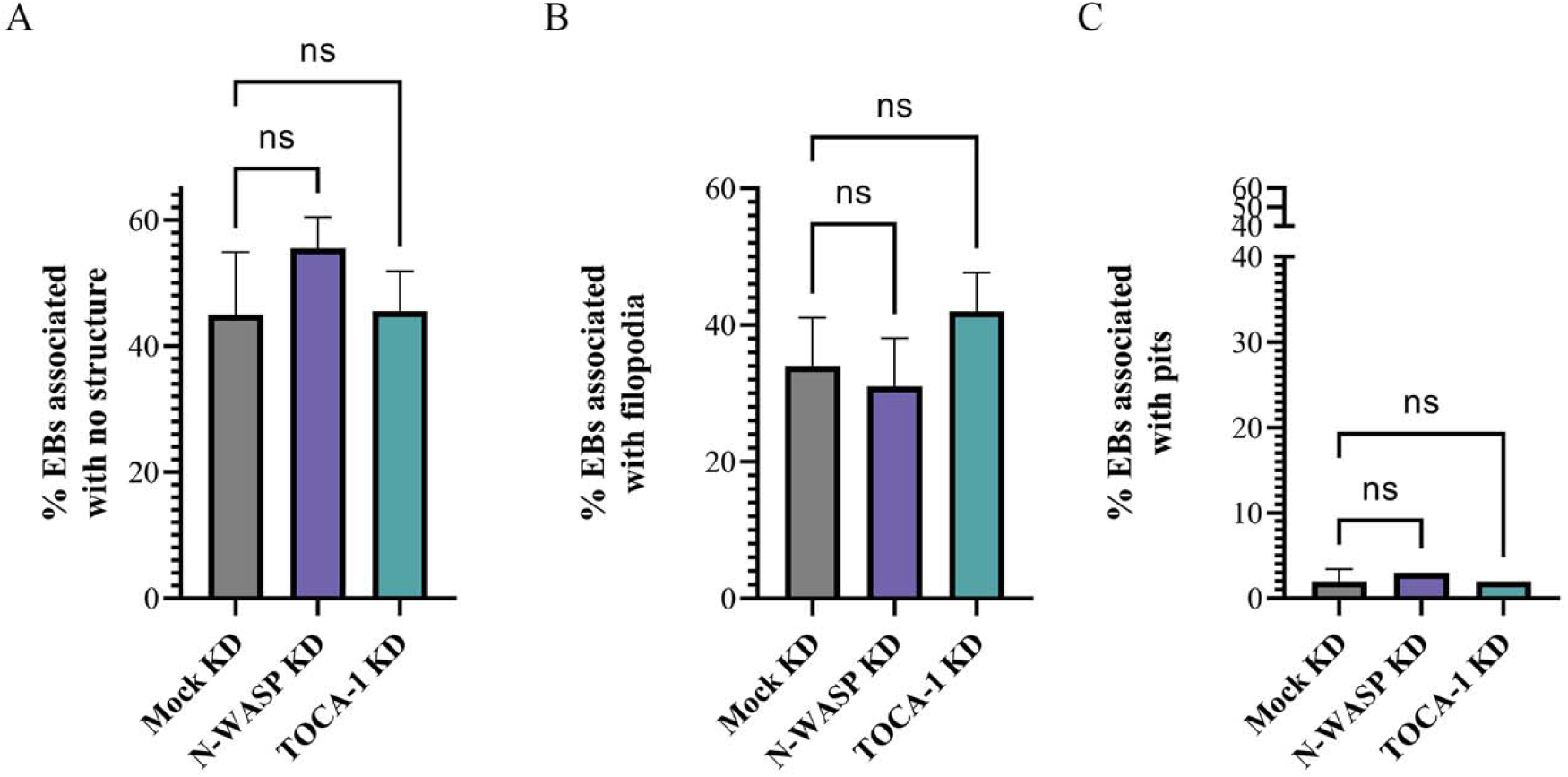
N-WASP and TOCA-1 KD do not impact formation of other structures. Quantification of EBs associated with **A.** no structure, **B.** filopodia or **C.** pits. 100 EBs per experiment were assessed from two separate experiments, in which images were blinded and categorized based on structure association. EBs in each category were compared to total EBs to determine the percentage associated with structures.

**Supplemental Figure 2.**
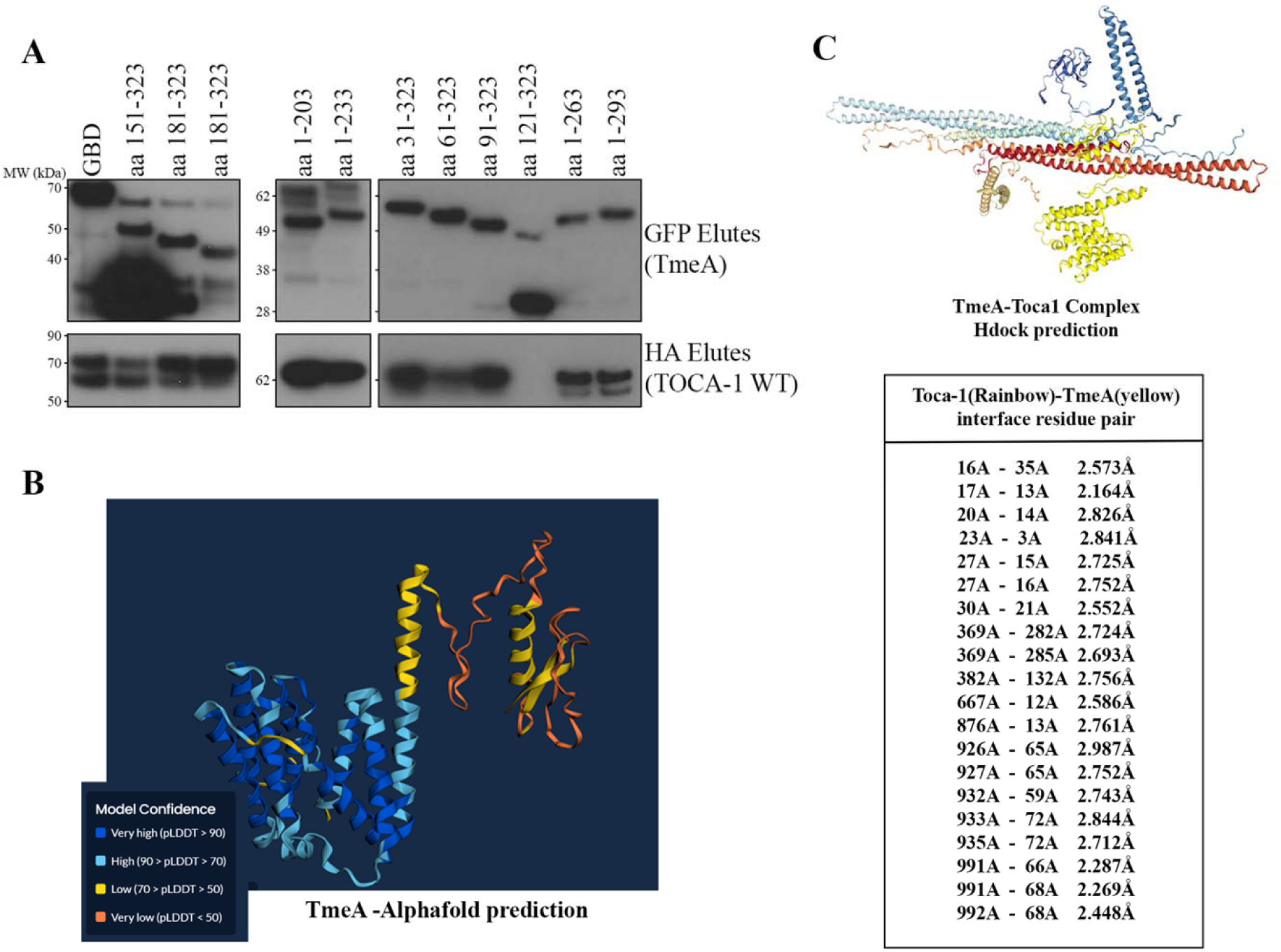
TmeA likely binds TOCA-1 at multiple binding sites. **A.** GFP-tagged TmeA truncations were co-transfected with HA-tagged TOCA-1 WT in HeLa cells. The GFP tagged proteins were immunoprecipitated and samples were probed with anti-GFP or anti-HA antibodies. **B.** Alphafold modeling predicting tertiary structure of TmeA. **C.** Binding complex prediction of TmeA (yellow, ligand) and TOCA-1 (rainbow, receptor). Confidence score is 0.9777, thus TmeA and TOCA-1 are very likely to bind. Residue pairs with predicted distance < 3Å are noted.

## Notes

### Competing Interest Statement

The authors have declared no competing interest.

